# metaFlye: scalable long-read metagenome assembly using repeat graphs

**DOI:** 10.1101/637637

**Authors:** Mikhail Kolmogorov, Mikhail Rayko, Jeffrey Yuan, Evgeny Polevikov, Pavel Pevzner

## Abstract

Long-read sequencing technologies substantially improved assemblies of many isolate bacterial genomes as compared to fragmented assemblies produced with short-read technologies. However, assembling complex metagenomic datasets remains a challenge even for the state-of-the-art long-read assemblers. To address this gap, we present the metaFlye assembler and demonstrate that it generates highly contiguous and accurate metagenome assemblies. In contrast to short-read metagenomics assemblers that typically fail to reconstruct full-length 16S RNA genes, metaFlye captures many 16S RNA genes within long contigs, thus providing new opportunities for analyzing the microbial “dark matter of life”. We also demonstrate that long-read metagenome assemblers significantly improve full-length plasmid and virus reconstruction as compared to short-read assemblers and reveal many novel plasmids and viruses.

## Introduction

Bacterial genome assemblies of long Single Molecule Sequencing reads (generated by Pacific Biosciences or Oxford Nanopore sequencers) substantially improved the contiguity of assembled genomes as compared to short-read assemblies (Phillippy, 2017, Jain et al., 2018, Schmid et al., 2018). In contrast, early long-read metagenomic studies reported lower yield and reduced read length compared to isolate bacterial assemblies, making it difficult to generate high-quality assemblies and suggesting that sample preparation protocols have to be optimized to utilize long reads in metagenomic studies (Tsai et al., 2016, Driscoll et al., 2017). However, the recent improvements in high molecular weight DNA extraction techniques allow one to sequence complex metagenomes with deep coverage and increased read length (Moss and Bhatt. 2018, Bertrand et al., 2018, Somerville et al., 2018, Nicholls et al., 2019). These improved protocols have already been used for sequencing complex bacterial communities (Bickhart et al., 2018, Stewart et al., 2018).

Although some long-read assemblers (Chin et al, 2016, Li, 2016, Koren et al., 2017, Kamath et al., 2017, Kolmogorov et al., 2019, Ruan and Li, 2019) have been applied to metagenomic datasets, none of them was specifically designed for metagenome assembly. This is unfortunate since long-read metagenomic assemblies have a potential to greatly increase the contiguity of short-read assemblies and address their inherent limitations, such as strain resolution (Goltsman et al., 2018), detection of horizontal gene transfer (Guo et al., 2015), and sequencing of novel plasmids and viruses (Arredondo-Alonso et al., 2017, Paez-Espino et al., 2016).

Metagenomic assembly presents additional computational challenges compared to the assembly of isolates due to highly non-uniform coverage of the species/strains comprising the sample, the presence of long intra-genomic and inter-genomic repeats (Li et al., 2015, Nurk et al., 2017), and difficulties in plasmid and virus reconstructions (Antipov et al., 2019, Wick and Holt, 2019). We recently developed a fast, long-read genome assembler Flye and showed that it produces accurate and contiguous assemblies (Kolmogorov et al., 2019). Wick and Holt, 2019 benchmarked Flye on various bacterial datasets and demonstrated that it improves on the state-of-the-art long-read assemblers. Below we describe a fast, long-read metagenome assembler metaFlye, benchmark it on both mock and real bacterial communities, and demonstrate that it generates high quality assemblies.

## Results

The Flye algorithm first attempts to approximate the set of *genomic k*-mers (*k*-mers that appear in the genome) by selecting *solid k*-mers (high-frequency *k*-mers in the read-set). It further uses solid *k*-mers to efficiently detect overlapping reads, and greedily combines overlapping reads into *disjointigs* (Kolmogorov et al., 2019). This approach excludes most erroneous *k*-mers (that appear in reads but not in the genome) from consideration and reduces the memory footprint of the *k*-mer index. However, in a metagenome setting, this approach would favor high-abundance species, while low-abundance species will have a reduced number of solid *k*-mers (if any), and thus will fail to be assembled. To address this limitation, we introduce a new approach to solid *k*-mer selection, which combines global *k*-mer counting with analyzing local *k*-mer distributions (see Methods).

In difference from contigs (that are expected to represent contiguous segments of a genome), error-prone disjointigs represent arbitrary paths in the assembly graph that can be generated much faster than contigs. To fix potential misassemblies within disjointigs, Flye constructs the *repeat graph* from disjointigs by collapsing each family of long repeats into a single path in the graph (Kolmogorov et al., 2019). It further classifies edges in the repeat graph into *unique* and *repetitive* and simplifies the graph by untangling most repeat edges using *bridging reads*.

Flye classifies an edge in the repeat graph as repetitive if it has high coverage or if the reads that are traversing this edge induce diverging paths in the graph. Although the coverage-based criteria works well for isolate genome assembly, it is not applicable to metagenomic assembly. Further, the path-based criteria might fail to identify edges that belong to complex mosaic repeats. To address these complications, we developed a new repeat classification algorithm that reliably detects repeat edges in the metagenomic assembly graph in an iterative manner (see Methods).

In addition to the challenges of assembling bacterial chromosomes, there are additional difficulties in assembling short plasmids that are typically covered only by a small number of reads. We show that such plasmids often remain undetected by existing assemblers and describe an algorithm that recovers unassembled plasmids from long-read sequencing data.

We benchmarked metaFlye, Canu (Koren et al., 2017), miniasm (Li, 2016) and wtdbg2 (Ruan and Li, 2019) using three Pacific Biosciences (PacBio) and Oxford Nanopore Technology (ONT) mock metagenome datasets, for which closely related reference genomes are available. We also ran the FALCON assembler (Chin et al, 2016) on the PacBio datasets, but not on the ONT datasets (since FALCON requires PacBio-specific information as input). For each mock metagenome, we used metaQUAST (Mikheenko et al., 2018) to evaluate the statistics of the combined references (Table 1) as well as to compute the separate statistics for each species present in the sample (Figure 1). Figure 2 additionally shows NGAx plots for all datasets. Because miniasm outputs contigs with a high per-nucleotide error rate, we performed a round of contig polishing using Racon (Vaser et al., 2017) to generate more accurate miniasm assemblies (see Methods).

**Table 1.** Assembly statistics and benchmarks for the HMP, ZymoEven, and ZymoLog datasets. For the HMP dataset, we used 19 bacterial references with sufficient read coverage and excluded three references with low coverage for assembly quality evaluation with metaQUAST. All reference genomes were used for assembly quality evaluation in the case of the ZymoEven datasets. For the ZymoLog GridION dataset, we used only four reference genomes that had read coverages above 3x (*L. monocytogenes*, *P. aeruginosa*, *B. subtilis*, and *S. cerevisiae*) for assembly quality evaluation. Similarly, five bacteria and one yeast references were used in ZymoLog PromethION analysis. Two yeast genomes (*S. cerevisiae* and *C. neoformans*) were excluded from the misassembly analysis in all Zymo datasets because of the many apparent differences between the reference and the assembled strains. Statistics were computed with metaQUAST 5.0 with the sequence identity threshold set to 90% and the minimum contig length set to 5 Kb. All tools were benchmarked on a computational node with 52 Intel Xeon 8164 CPUs. Reference coverage is the percentage of the reference genome covered by assembled contigs. NGA50 is the NG50 statistic computed for contigs that are broken at their misassembly breakpoints.

| Dataset description | Assembler | Total length | Contigs | Reference coverage | NGA50 | Mis-assemblies | CPU hours |
| --- | --- | --- | --- | --- | --- | --- | --- |
| HMP<br>6.8 Gb PacBio,<br>19 bacterial<br>ref. | metaFlye | 66.2 Mb | 85 | 99.8% | 1.77 Mb | 67 | 45 |
|  | Canu | 68.2 Mb | 180 | 99.7% | 1.82 Mb | 122 | 756 |
|  | FALCON | 60.0 Mb | 388 | 90.3% | 0.76 Mb | 116 | 150 |
|  | miniasm + Racon | 66.5 Mb | 95 | 99.6% | 1.48 Mb | 74 | 11 |
|  | wtdbg2 | 65.5 Mb | 187 | 98.7% | 0.66 Mb | 104 | 4 |
| ZymoEven<br>GridION<br>14 Gb ONT,<br>8 bacterial & 2<br>yeast ref. | metaFlye | 65.5 Mb | 637 | 94.6% | 526 Kb | 29 | 90 |
|  | Canu | 65.7 Mb | 807 | 95.6% | 272 Kb | 36 | 4,590 |
|  | miniasm + Racon | 51.9 Mb | 998 | 80.4% | 83 Kb | 26 | 67 |
|  | wtdbg2 | 54.2 Mb | 1101 | 75.9% | 75 Kb | 11 | 5 |
| ZymoLog<br>GridION<br>16 Gb ONT,<br>3 bacterial & 1<br>yeast ref. | metaFlye | 23.6 Mb | 363 | 75.2% | 407 Kb | 10 | 112 |
|  | Canu | 26.9 Mb | 433 | 90.2% | 187 Kb | 57 | 38,800 |
|  | miniasm + Racon | 15.6 Mb | 122 | 56.5% | 209 Kb | 43 | 299 |
|  | wtdbg2 | 23.0 Mb | 668 | 56.6% | 14 Kb | 20 | 13 |
| ZymoEven<br>PromethION<br>146 Gb ONT<br>8 bacterial & 2<br>yeast ref. | metaFlye | 70.3 Mb | 527 | 94.6% | 647 Kb | 11 | 1,000 |
|  | wtdgb2 | 25.7 Mb | 313 | 39.4% | - | 30 | 12 |
| ZymoLog<br>PromethION<br>148 Gb ONT<br>5 bacterial & 1<br>yeast ref. | metaFlye | 38.4 Mb | 284 | 95.4% | 3 Mb | 34 | 4,500 |
|  | wtdgb2 | 17.3 Mb | 385 | 32.8% | - | 31 | 16 |

**Figure 1.**
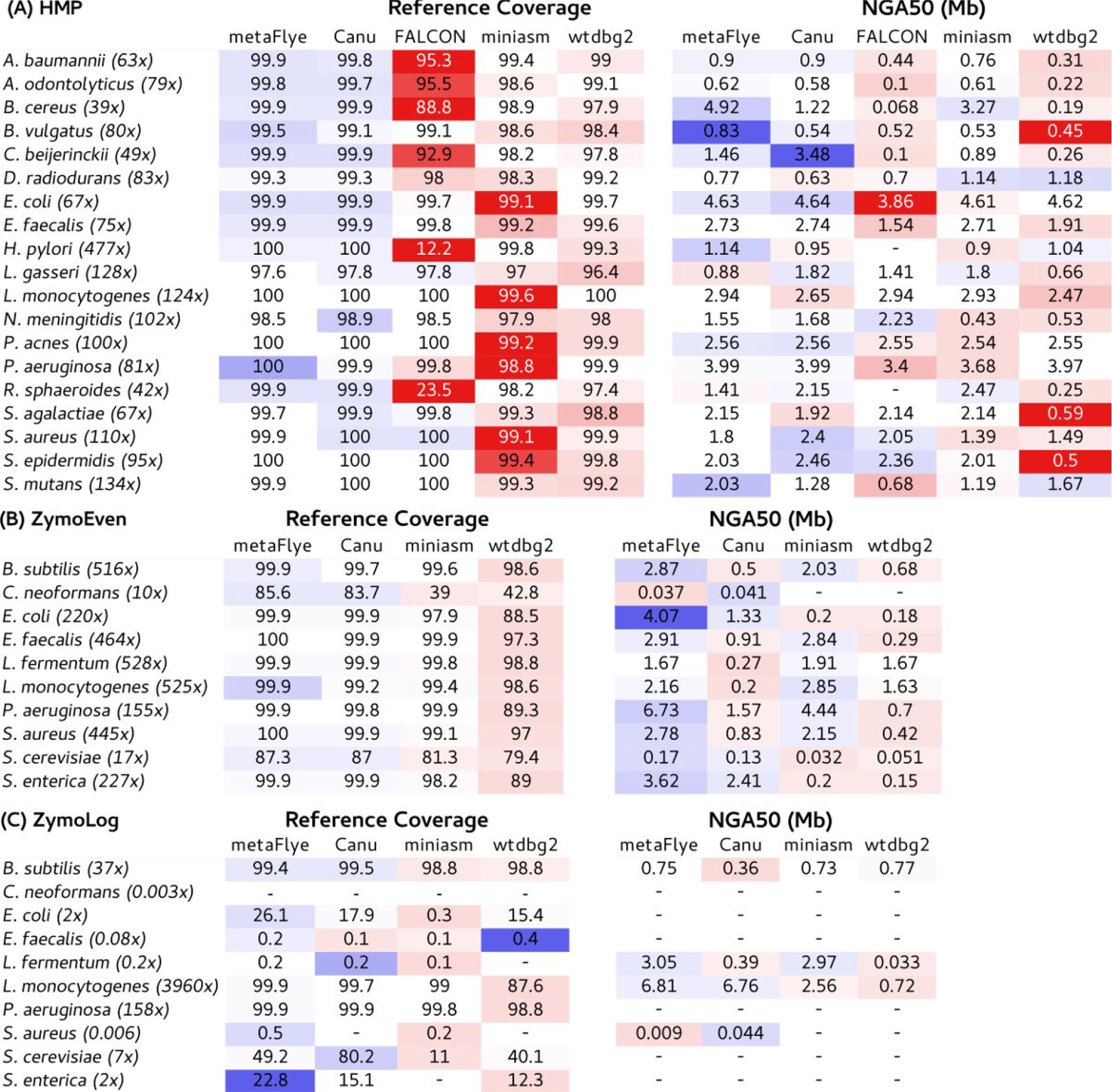
Per-species statistics for the HMP, ZymoEven GridON, and ZymoLog GridON datasets. Reference coverage and NGA50 statistics were computed using metaQUAST. The read coverage for each species are given in the brackets after the species name. NGA50 values are not reported for assemblies with reference coverage below 50%.

**Figure 2.**
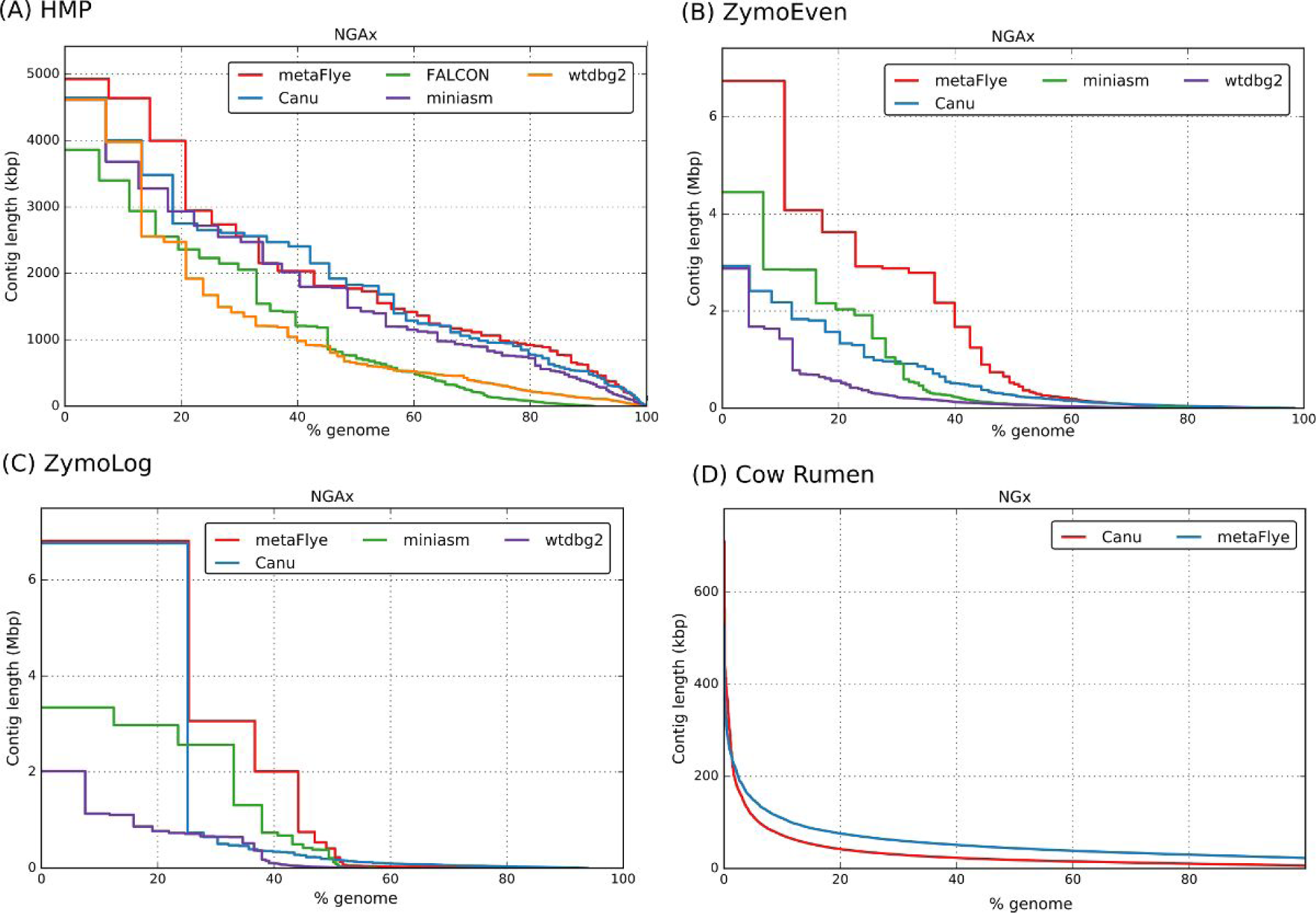
NGAx / NGx plots for different datasets generated using metaQUAST. NGA statistics were computed for the datasets with available references (A-C). For the cow rumen dataset (D), the NG plot is given with the genome size set to 1 Gb.

The HMP mock dataset represents a human mock metagenome formed by 22 bacteria with known reference genomes sequenced using PacBio reads (total length 6.8 Gb and N50 = 6.7 kb). Nineteen of these bacteria have read coverages ranging from 39x (*B. cereus*) to 477x (*H. pylori*). Since the remaining three genomes (*M. smithii, C. albicans*, and *S. pneumoniae*) have low coverage (below 1x), they were excluded from further analysis.

The metaFlye, Canu and miniasm assemblies resulted in high reference coverage (ranging from 99.6% for miniasm to 99.8% for metaFlye) and NGA50 (ranging from 1.48 Mb for miniasm to 1.82 Mb for Canu). The number of mis-assemblies varied from 67 for metaFlye to 122 for Canu. The wtdbg2 and FALCON assemblies had reduced reference coverage (98.6% and 89.9%, respectively) and lower contiguity (NGA50 = 0.66 Mb and 0.76 Mb, respectively). The reduced coverage and contiguity were mainly associated with bacteria with abundances that significantly deviated from the median dataset coverage (*B. cereus, R. shaeroides, C. beijerinckii* and *H. pylori;* see Figure 1), highlighting the challenge of assembling metagenomics datasets with uneven species abundance.

metaFlye assembled all 14 known plasmids that have been previously identified in the HMP dataset (Antipov et al., 2019). Miniasm, Canu, FALCON and wtdbg2 missed one, two, four, and four plasmids, respectively. Most of the missed plasmids were shorter than 5 Kb and were fully covered by a single read, illustrating additional complications in reconstructing short plasmids. Overall, plasmid reconstruction using long reads showed substantial improvement over short-read metagenome assemblers: metaplasmidSPAdes short-read plasmid assembler reconstructed only seven out of the 14 plasmids from the same sample (Antipov et al., 2019).

We then evaluated the assemblies of the mock datasets from the ZymoBIOMICS Microbial Community Standards, generated using ONT reads with an N50 of ~5 Kb (Nicholls et al., 2019). The *ZymoEven* mock community consists of eight bacteria with abundance ≅12% and two yeast species with abundance ≅2%. The *ZymoLog* dataset represents the same microbial community with abundances distributed as a log scale from 89.1% (*Listeria monocytogenes*) to 0.000089% (*Staphylococcus aureus*). Each of the two communities were sequenced using GridION (total read lengths of 14 Gb and 16 Gb for the ZymoEven and ZymoLog datasets, respectively) and PromethION (total read lengths of 146 Gb and 148 Gb for the ZymoEven and ZymoLog datasets, respectively). Since the provided reference assemblies of two yeast species (*S. cerevisiae* and *C. neoformans*) were highly fragmented, we substituted them with the closest complete reference strains from NCBI (YJM1307 and JEC21, respectively). Because of the structural differences between the reference and the assembled strains, we ignored misassemblies from the yeast genomes in the total count of misassemblies.

The Canu and metaFlye assemblies of the ZymoEven GridION dataset covered 95.6% and 94.6% of the combined reference length (in contigs of length 5 kb and higher), and significantly improved over miniasm and wtdbg2 assemblies (80% and 76%, respectively). metaFlye showed better contiguity than Canu (NGA50 = 526 kb and 272 kb, respectively). The number of misassemblies varied from 11 for wtdbg2 to 36 for Canu. Figure 1 illustrates that metaFlye and miniasm produced very similar assemblies of most of the bacterial species; however, miniasm produced more fragmented assemblies of the yeast species.

The ZymoLog GridION dataset has only four species with read coverage above 3x: *L. monocytogenes* (3960x), *P. aeruginosa* (158x), *B. subtilis* (38x) and *S. cerevisiae* (7x). metaFlye reconstructed over 99% of the three most abundant bacteria and 49% of *S.cerevisiae*. Miniasm assembled a smaller fraction of *S.cerevisiae* (10%), and wtdbg2 generated a highly fragmented assembly of the abundant *L. monocytogenes.* Canu produced the best coverage of the *S.cerevisiae* genome (80%), however the assembly was highly fragmented. Overall, metaFlye showed the best contiguity, followed by miniasm and Canu (NGA50 = 407 Kb, 209 Kb and 187 Kb, respectively; see Figure 1). The number of misassemblies varied from 10 for metaFlye to 57 for Canu (Table 1).

metaFlye assembled PromethION runs of both ZymoEven and ZymoLog communities in 1,000 and 4,500 CPU hours, respectively (Table 1). In the ZymoEven dataset, all bacterial genomes but two were assembled into single circular contigs (assemblies of *L. monocytogenes* and *E. faecalis* resulted in three contigs since they share an unresolved repeat of length 35 kb). The contiguity of the *C. neoformans* and *C. cerevisiae* assemblies (NGA50) increased by a factor of 2 as compared to the GridION assembly. The metaFlye assembly of the ZymoLog community from PromethION reads significantly improved over its assembly of the community from GridION reads: the cumulative reference coverage of all species increased from 38% to 58%. In particular, *S.cerevisiae* coverage increased from 49% to 87% and the previously unassembled *E. coli* and *S. enterica* genomes had over 99% coverage in the PromethION assembly. This improvement in the reconstruction of underrepresented species highlights the benefits of generating deep coverage datasets for long-read metagenome sequencing.

Since the Canu running time on the Zymo PromethION datasets was estimated as 50,000+ CPU hours, it was impractical to run it on the available hardware. Miniasm requires all pairwise-read alignments to be computed first using minimap2. The size of such alignment (in the PAF format) for the GridION ZymoEven dataset was ~200 Gb. Since the expected size of the alignments for the PromethION dataset is ~20 Tb (the number of alignments is quadratic in the number of reads), it was impractical to run miniasm on the available hardware. The wtdbg2 assembly of the ZymoEven PromethION dataset was much shorter than its GridION assembly (26 Mb vs 54.2 Mb, respectively), which might be a result of read subsampling procedures implemented in this assembler. Similarly, the ZymoLog assembly size was reduced from 23.4 Mb for GridION to 17.3 Mb for PromethION.

In addition to the mock metagenomic datasets, we assembled the cow rumen dataset consisting of PacBio reads (total read length 52.2 Gb with N50 ~9 Kb) and compared metaFlye assembly against Canu assembly generated in the original study (Bickhart et al., 2018). Both assemblies were polished using short reads with two rounds of the Pilon polishing procedure in the indel correction mode (Walker et al., 2014). The metaFlye and Canu assemblies had total lengths of 1,260 Mb and 1,035 Mb in contigs longer than 5 Kb, respectively. The metaFlye assembly was also more contiguous (Figure 2): the NG50 was 44 Kb for metaFlye and 19 Kb for Canu, for a hypothetical metagenome size of 1 Gb (NGAalignment statistics were not computed since the reference genomes were unknown). Prodigal (Hyatt et al., 2010) predicted 1,431,527 full (38,376 partial) genes in the metaFlye assembly, and 1,191,681 full (95,679 partial) genes in the Canu assembly.

We then used Barrnap (https://github.com/tseemann/barrnap) to identify 581 and 422 full-length 16S rRNA genes in metaFlye and Canu assemblies, respectively. We further performed Operational Taxonomic Unit (OTU) clustering of these genes at 95% identity using QIIME2 (Bolyen et al., 2018) to reveal the fine-grained taxonomic composition of the microbial community. The clustering resulted into 377 and 218 OTUs for metaFlye and Canu assemblies, respectively (all but one OTUs from Canu assembly were also recovered by metaFlye). Taxonomic assignment against SILVA132 full-length rRNA database (Quast et al., 2012) revealed 65 and 34 unassigned OTUs (with best match against the SILVA132 database below 95% percent identity) in metaFlye and Canu assemblies, respectively. This analysis reveals the great potential of long-read metagenomic assemblers to analyze the microbial “dark matter of life” (Lloyd et al., 2018). In contrast to short-read assemblers that typically fail to reconstruct full-length 16S RNAs (due to collapsing their multiple copies and further assembly fragmentation), long-read assemblers capture many 16S RNA genes within long contigs (up to 374 kb long in the metaFlye assembly).

To identify Antibiotic Resistance Genes (ARGs), we predicted ORFs with Prodigal in the metagenomic mode, aligned them using blastn against the ResFinder ARG database (Zankari et al., 2012), and retained hits with at least 95% nucleotide sequence identity and at least 90% ARG sequence coverage. This procedure resulted in 46 and 104 putative ARGs in metaFlye and Canu assemblies, respectively. To rule out a possibility that some of them are caused by duplicated regions in assemblies, we clustered the identified ARGs based on their *k*-mer content using Mash (Ondov et al. 2016) and classified two ARGs as duplicates if their *k*-mer compositions differ by less than 1%. After this clustering, we revealed 26 and 23 potential unique ARGs in metaFlye and Canu assemblies, respectively (16 of them were found in both metaFlye and Canu assemblies).

plasmidVerify (Antipov et al., 2019) and VirSorter (Roux et al, 2015) identified 52 putative plasmids and 37 viruses among circular contigs in the metaFlye assembly. Among them, seven plasmids and two viruses were identified only using the plasmid detection algorithm aimed at short plasmids and described in Methods section. All but three of the identified plasmids and viruses did not have significant BLAST matches against the NCBI database, thus potentially representing novel plasmids/viruses.

plasmidVerify and VirSorter identified fewer putative plasmids among circular contigs in Canu assembly of the cow rumen microbiome (39 versus 52) but more viruses (87 versus 37) as compared to the metaFlye assembly. Interestingly, there is little overlap between plasmids (only 8) and viruses (only 10) identified in metaFlye and Canu assemblies, suggesting that there may be a synergy between these two tools with respect to plasmid and virus assembly.

## Discussion

Although long-read metagenomics is a promising direction for untangling complex bacterial communities, it faces difficult algorithmic challenges. We developed the first long-read metagenomic assembler metaFlye and benchmarked it using both mock and real metagenomic communities. Most long-read assemblers generated assemblies with a high reference coverage of the HMP mock dataset, with metaFlye, Canu and miniasm assemblies being the most contiguous. However miniasm and wtdbg2 had difficulties in assembling species with large deviations in coverage in the Zymo datasets, and the Canu assemblies showed reduced contiguity and an increased number of assembly errors. With respect to the running time, metaFlye was 10 to 300-fold faster than Canu on the various datasets. Only metaFlye and wtdgb2 were able to scale to the 150 Gb PromethION runs, but wtdbg2 assemblies were surprisingly incomplete and fragmented, compared to the corresponding GridION assemblies.

Our analysis of the cow rumen dataset revealed that long-read assemblers greatly improve on short-read assemblers with respect to full-length sequencing of 16S RNA genes, plasmids and viruses, metaFlye and Canu assemblies of this dataset confirmed the trend that we observed with the mock metagenomes, with the metaFlye assembly being more contiguous (in terms of the NG50 statistics) than the one produced by Canu. Since the number of assembly errors is not known, It remains unclear what is the gap between the NGA50 and NG50 statistics for both Canu and metaFlye (note that Canu assemblies of all mock datasets had the largest numbers of misassemblies among all benchmarked tools).

Although metaFlye is currently limited to assembling long reads only, we plan to extend it to assembling hybrid datasets that combine long and short reads. The existing hybrid assemblers, such as hybridSPAdes (Antipov et al., 2016) and Unicycler (Wick et al., 2017), first assemble short reads using SPAdes or metaSPAdes and further scaffold the resulting contigs using *individual* long reads. As a result, they do not fully utilize long-read assemblies. metaFlye enables an alternative approach based on (i) assembling long reads into metaFlye contigs, (ii) assembling short reads into metaSPAdes contigs, and (iii) combining metaFlye and metaSPAdes contigs and assembling them together using Flye.

## Methods

### Generating assemblies

metaFlye was run using the following additional options for the mock metagenome datasets: “--*meta* --*plasmids*”. While assembling the cow rumen dataset, we found that 13% of PacBio reads contained more than one PacBio subread (reads with multiple polymerase passes). To efficiently split those “chimeric” reads, we developed a small program called pbclip (https://github.com/fenderglass/pbclip) and applied it to the PacBio data before running metaFlye. Minimum overlap parameter for metaFlye was manually set to 2 kb for the cow rumen assembly. Miniasm was run using their default parameters for all datasets. Canu was run using parameters recommended for metagenome assembly: “*corOutCoverage*=*10000 corMhapSensitivity*=*high corMinCoverage*=*0 redMemory*=*32 oeaMemory*=*32 batMemory*=*200*”. Wtdbg2 was run using the default parameters for the HMP dataset. However, since the Zymo datasets had higher read coverage as well as low-abundant species, we increased the *k*-mer frequency coverage range by using “--*node*-*max 1000* -*e 2*” as suggested by the developers. This resulted in a significant increase in the total length of the assembly as compared to the default settings (from 28 Mb to 55 Mb for the ZymoEven dataset, and from 12.6 Mb to 23.4 Mb for the ZymoLog dataset). FALCON was run using a configuration file recommended for bacterial assemblies.

### Solid *k*-mer selection in metagenome assemblies

The Flye algorithm (Kolmogorov et al., 2019) selects solid *k*-mers as follows (the typical *k*-mer size is 15 or 17 nucleotides for PacBio and ONT reads). In the first pass through all reads, the algorithm counts frequencies of *k*-mer *hashes* using a fixed-size array of counters. In the second pass, *k*-mers with pre-computed frequency higher than a threshold (typically equal to 2 or 3) are counted using a cuckoo hash table (Li et al., 2014). Given the computed *k*-mer frequency table and an estimated genome size |*G*|, the algorithm selects the |*G*| most frequent *k*-mers, and sets a frequency threshold *t* as the minimum frequency among the selected *k*-mers. The selected threshold *t* separates solid *k*-mers (that are indexed) from erroneous ones (that are discarded).

This strategy typically results in a relatively small misclassification rate; e.g., in a typical isolate bacterial project only ~5% of unique *genomic k*-mers (true *k*-mers from the genome) are missing from the set of solid *k*-mers and only ~10% of unique solid *k*-mers represent non-genomic *k*-mers. However, although it works well in genomic assemblies, it is not suitable for metagenomic assemblies, because there is no frequency threshold that robustly separates genomic from non-genomic *k*-mers (due to the uneven species coverage). Below, we describe an alternative strategy for solid *k*-mer selection and benchmark it using both isolate and metagenome datasets.

Similarly to the uniform coverage mode in Flye, metaFlye also starts with counting *k*-mers in all reads. Although high-frequency *k*-mers are still expected to represent genomic *k*-mers, non-genomic *k*-mers arising from reads in highly abundant species often outnumber genomic *k*-mers from rare species. Given a per-nucleotide error rate *ɛ* in reads, we estimate the probability of a *k*-mer in a read to be error-free as *E* = *e*^−*kɛ*^, under a Poisson error distribution model. Thus, the expected number of solid *k*-mers in a read is *E* * |*read*|. For each read, metaFlye selects a frequency threshold *f* so as there are at least *E* * |*read*| *k*-mers in this read with frequency at least *f* and indexes *k*-mers above this threshold using a hash table. Similarly to other *k*-mer counting/indexing tools, metaFlye keeps the canonical representation of each *k*-mer, which is defined as the lexicographical minimum of the forward and reverse-complement of the *k*-mer.

We evaluated the uniform and metagenome *k*-mer selection modes using two bacterial datasets, for which true *k*-mers were extracted from the available references. The first set of PacBio reads from an *E. coli* isolate (at 50x coverage) contains 254.2M (million) *k*-mers, out of which 56.7M (22%) are genomic. In the uniform *k*-mer selection mode, Flye indexed 55.3M genomic *k*-mers (97% of all genomic *k*-mers) and 5.0M non-genomic (erroneous) *k*-mers. In the metagenome selection mode, metaFlye indexed 50.3M genomic *k*-mers (89%) and 22M non-genomic *k*-mers.

We further used the HMP dataset (described above) to evaluate the *k*-mer selection in a metagenome setting. We focused on the two least abundant genomes in the mixture – *B. cereus* and R. *sphaeroides* – which had coverage 2-fold below the median coverage. These two bacteria contributed to 83M genomic *k*-mers in the reads. In the uniform coverage mode, Flye selected only 33.2M (40%) of their genomic *k*-mers. In contrast, metaFlye selected 71M (86%) of genomic *k*-mers in the metagenome coverage mode.

### Identifying repeats in the metagenome assembly graph

metaFlye classifies each edge of the metagenome assembly graph as *unique* (its sequence appears only once in a single genome) or *repetitive* (the edge sequence appears multiple times in a single genome or is shared by multiple genomes). Flye uses this classification to identify *bridging reads* (that start and end at different unique edges) and resolves repeats using bridging reads (Kolmogorov et al., 2019). Thus, the contiguity of Flye assemblies critically depends on its ability to correctly classify unique and repetitive edges of the assembly graph.

The Flye algorithm first aligns all reads to the assembly graph, computes the mean coverage of each edge and represents all reads as *read-paths* (paths in the assembly graph). Afterwards, it captures the lion’s share of the repeat edges by simply classifying all high-coverage edges (with coverage exceeding the mean coverage by a factor of 1.75) as repetitive. However, since there are possible variations in coverage along the genome, this procedure mis-classifies some repetitive edges as unique. To improve the classification of such edges, Flye additionally checks whether all read-paths through a unique edge continue into a single *successor* edge (a similar test is done for *predecessor* edges). If there are multiple successors or predecessors, the edge is re-classified as repetitive.

Although this approach works well in genomic assemblies, it is not suitable for metagenomic assemblies since the edge coverage is not a reliable predictor of the edge multiplicity. Without the coverage test, the read-paths criteria might fail to identify repetitive edges that belong to mosaic repeats, since it only checks one immediate predecessor and successor of each edge (Figure 3). To address this pitfall, we substitute the “diverged read-paths” approach in Flye by the “repeat detection” approach in metaFlye (described below) to identify repeat edges in the metagenome assembly graph without using coverage information.

**Figure 3.**
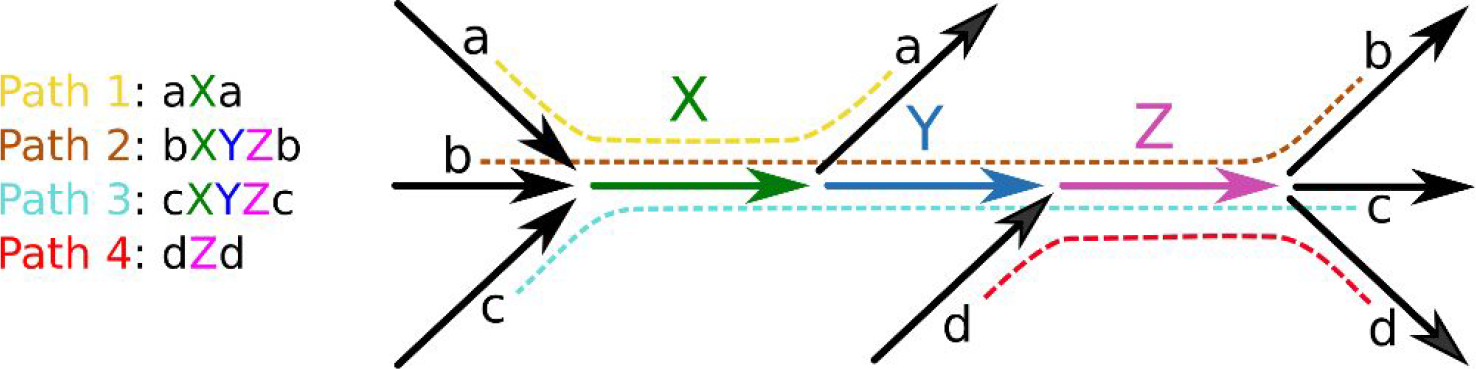
An example of a mosaic repeat. The subgraph of an assembly graph is formed by four distinct genome sub-paths. Edges are shown in color (for repeats of multiplicity 2 or 3), or in black (for unique edges of multiplicity 1). Although the edge *Y* is a part of the mosaic repeat, Flye may classify it as a unique edge since it has a single predecessor (*X*) and a single successor (*Z*). The metaFlye repeat detection algorithm will classify *X* and *Z* as repetitive on the first iteration (since they have three predecessors / successors). On the second iteration, *Y* will be classified as a repeat, since there exist reads that start at *Y* and continue into multiple predecessors / successors of *X* and *Z*, thus revealing that *Y* is a repeat.

Initially, all edges in the assembly graph are labeled as unique. The algorithm iterates through all edges and may change their classification into repetitive as described below. Thus, at each intermediate iteration, the assembly graph may contain both unique and repetitive edges.

Given a read-path through an edge *e*, metaFlye defines the next *unique* edge in this path as a *successor* of *e* (note that the original algorithm considers *any* edge as a successor). A set of all read-paths through an edge defines either a single or multiple successors. To account for chimeric reads, metaFlye filters out successors that are supported by less than *MaxSucc* / *delta* reads, where *MaxSucc* is the number of reads for a successor with the highest support and *delta* is a threshold (the default value of *delta*=5). If an edge has multiple successors or predecessors, it is classified as repetitive. The described test is performed iteratively on the entire set of edges, until no new edges are classified as repetitive.

Intuitively, in a mosaic repeat, the first iteration of the test will classify *some* of its edges as repetitive, but consecutive iterations extend the set of repeats (Figure 3). For a faster convergence of the algorithm, we traverse edges of the graph in the increasing order of their length, as short edges are more likely to be repetitive (two iterations are typically sufficient).

### Assembling short plasmids

We distinguish between short (shorter than the threshold *L* with the default value 10 kb)) and long plasmids (of length at least *L*). Sequencing of short plasmids is an important task since they represent a large fraction (~30%) of all plasmids in the RefSeq database. However, although existing long-read assemblers perform well in assembling long circular plasmids (longer than the typical read length), our benchmarking revealed that they often miss short plasmids. Paradoxically, the longer the reads, the more plasmids remain unassembled.

To assemble short plasmids, metaFlye first aligns all reads to the assembled contigs and then extracts unaligned reads (reads with an aligned fraction below 50%). It further extracts single unaligned reads and pairs of unaligned reads that assemble into circular sequences. metaFlye focuses on single reads and pairs of reads because short plasmids are typically fully covered by a single read or a pair of reads.

Given the set of unaligned reads, metaFlye constructs a set of short *cyclocontigs* by first selecting all self-overlapping reads, i.e., reads that have overlapping prefix and suffix. To further extend the set of cyclocontigs, it considers all pairs of reads and selects pairs (*A, B*) such that *B* overlaps *A* and *A* overlaps *B*. The collection of the constructed (unpolished) cyclocontigs may contain duplicates that represent the same circular plasmid. To extract unique sequences from the collection, metaFlye again performs an all-vs-all alignment of all the constructed cyclocontigs, finds similar ones and clusters them so that each cluster represents a unique circular sequence. metaFlye filters out single-read clusters (which are likely to represent artifacts from the extraction of unaligned reads). It then selects a representative for each cluster, polishes the representation using all reads contributing to the cluster as described in Lin et al., 2016, and adds these sequences to the final assembly output.

### Software versions used

- Flye: 2.4.2
- Canu: 1.8
- FALCON: pb-falcon 0.2.5
- Miniasm: 0.3
- Wtdbg2: 2.3
- QUAST: 5.0.2

## Data availability

The described datasets are available from the corresponding locations:

- HMP mock dataset:
https://github.com/PacificBiosciences/DevNet/wiki/Human_Microbiome_Project_MockB_Shotgun
- Zymo datasets: https://github.com/LomanLab/mockcommunity
- Cow rumen dataset: NCBI SRA repository under Bioproject PRJNA507739

Assemblies and metaQUAST evaluations used in this study are available at:
https://doi.org/10.5281/zenodo.2801953

## Code availability

metaFlye is freely available as a part of the Flye package at: https://github.com/fenderglass/Flye. The pbclip tool for PacBio subreads splitting is available from: https://github.com/fenderglass/pbclip.

## Acknowledgements

We are grateful to Derek Bickhart and Tim Smith for sharing the cow rumen dataset prior to journal publication.

